# SWITCHES: Searchable Web Interface for Topologies of CHEmical Switches

**DOI:** 10.1101/2020.08.04.232371

**Authors:** G.V. HarshaRani, S. Moza, N. Ramakrishnan, U.S. Bhalla

## Abstract

Bistable biochemical switches are key motifs in cellular state decisions and long-term storage of cellular ‘memory’. There are a few known biological switches that have been well characterized, however these examples are insufficient for systematic surveys of properties of these important systems. Here we present a resource of all possible bistable biochemical reaction networks with up to 6 reactions between 3 molecules, and 3 reactions between 4 molecules. Over 35,000 reaction topologies were constructed by identifying unique combinations of reactions between a fixed number of molecules. Then, these topologies were populated with rates within a biologically realistic range. The Searchable Web Interface for Topologies of CHEmical Switches (SWITCHES, https://switches.ncbs.res.in) provides a bistability and parameter analysis of over 7 million models from this systematic survey of chemical reaction space. This database will be useful for theoreticians interested in analyzing stability in chemical systems and also experimentalists for creating robust synthetic biological switches.

**Availability and Implementation:** Freely available on the web at https://switches.ncbs.res.in. Website implemented in PHP, MariaDB, Graphviz, and Apache, with all major browsers supported.

**Supplementary Information:** Not applicable.

## 1 INTRODUCTION

Biochemical switches are key motifs in cellular state decisions and long-term storage of cellular ‘memory’ (Angeli et al., 2004, Craciun et al., 2006, Markevich et al., 2004, Tyson et al., 2003). Such switches are bistable, that is, they have the property that they can exist stably in either of two states, with a third state acting as a transition or saddle point.

Two broad strategies to characterize bistable systems are the theory-based (e.g., Soranzo and Altafini, 2009, Craciun and Feinberg, 2006) and the catalog/database approach (Ramakrishnan and Bhalla, 2008). In the former, the graph structure of the network is used to determine whether the network has the potential for bistability irrespective of specific rate constants. The theoretical results underlying these methods are sometimes restrictive in their scope (e.g., the chemical reaction network theory methods used in (Soranzo and Altafini, 2009) do not permit zero concentrations at the steady states. Furthermore, these methods only guarantee multi-stationarity, not multi-stability and cannot make a conclusive determination of potential for bistability for some examples. In the current study we adopt the database approach. Here we catalog specific, fully defined chemical networks using linear algebra criteria for bistability, that is, two stable states and a saddle point.

## RESULTS

We previously conducted a systematic survey of chemical kinetic models to generate all possible reaction topologies up to a certain size, and assess their ability to form bistable switches (Ramakrishnan and Bhalla, 2008). The current database expands this survey to include over 7 million models sampled from 35,000 reaction topologies. encoded into a relational database form. 3561 of these topologies have at least one set of parameters that form a bistable model. There are over 33,000 bistable models, all of which have been encoded in SBML (Systems Biology Markup Language, Hucka et al., 2003). We provide interfaces for visualizing the chemical topology of each model in SBGN (Systems Biology Graphical Notation, Novère et al, 2009) format, and a way to navigate through relationships.

We generated chemical kinetic models using a small ‘alphabet’ of basic chemical reactions, assembling permutations of entries from this alphabet with permutations of molecules to participate in the reactions, eliminating isomorphisms, sampling parameters for each system, computing steady states, and then classifying them using linear algebraic criteria.

The database provides access to three key features of chemical space (**Fig 1A**). First, it organizes reaction systems by their topology and topology class. The topology of the chemical system is the configuration of how molecules react with each other, independent of the rate information. The topology class contains all the topologies with the same number of reactants and reactions. Second, it organizes the relationships between bistables. One of the key findings of (Ramakrishnan and Bhalla 2008) was that bistable chemical systems tend to be derived from simpler systems that are also bistable. In the SWITCHES database, we provide a graphical interface for users to navigate these relationships, both through a *family-tree* diagram of all switches and through individual records. Third, SWITCHES classifies all reaction systems in terms of their stability properties. For each reaction topology we sample 100 specific models using logarithmic Monte Carlo sampling (Ramakrishnan and Bhalla 2008). In addition, we use latin hypercube sampling (Moza, 2020, Deutsch and Deutsch, 2012) to sample more uniformly in the high-dimensional space of reaction-rates.

**Fig. 1.**
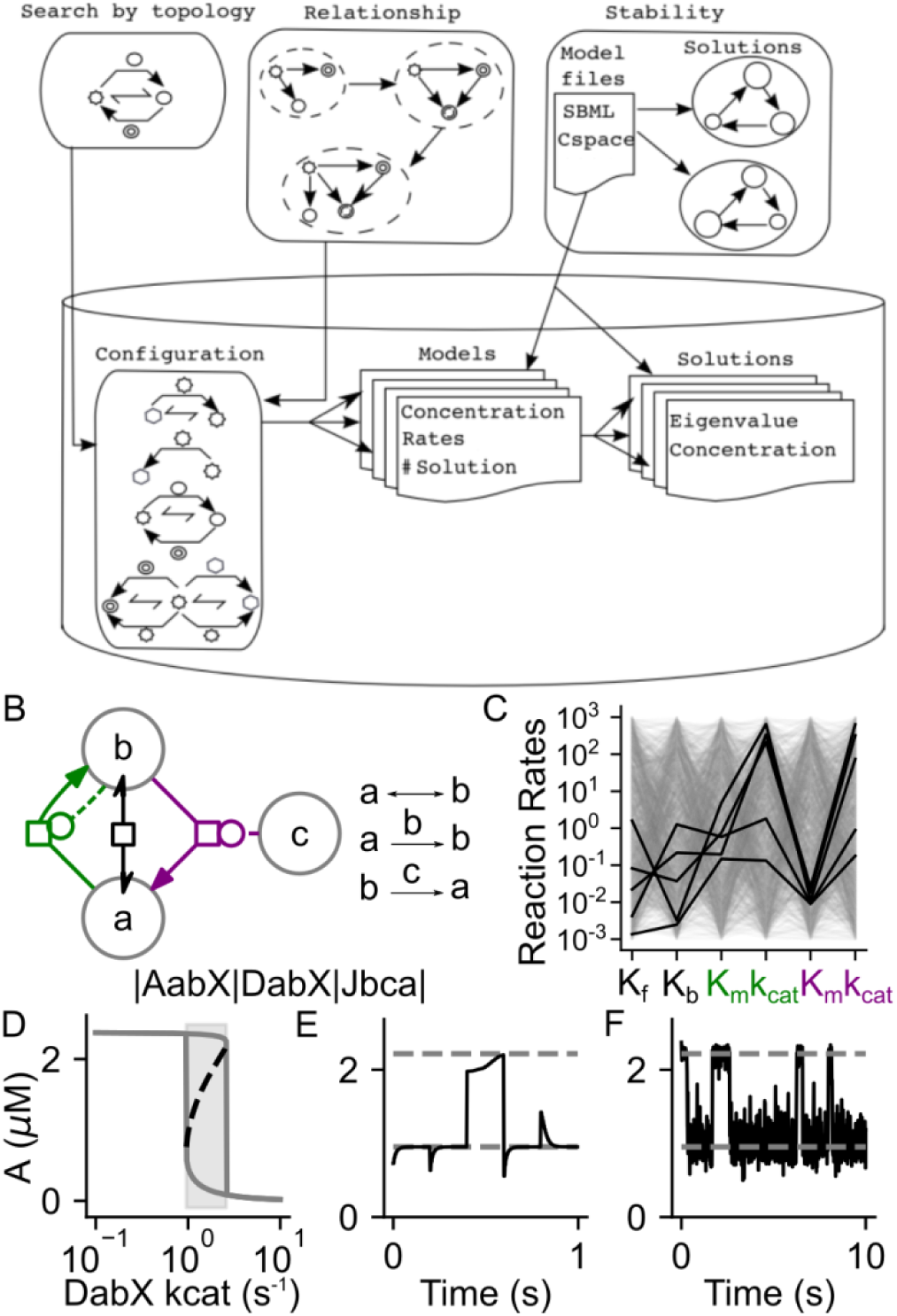
**A**. Interface and functional organisation of the SWITCHES database. The web interface supports searches by model topology, traversal through related models, and searches based on model solutions. Search results provide model files in SBML and condensed text (CSPACE) format, as well as eigenvalue analysis of solutions. The table structure maps to the reaction topology, the model parameters, and the model solutions respectively. **B-E** Characterizing the properties of an example bistable chemical reaction network. B. Chemical reaction network in the SBGN format on the left with corresponding chemical reactions on the right and CSPACE signature displayed below. C. A 1000 parameter sweep across three orders of magnitude for all rate parameters of this model using Latin Hypercube Sampling. Dark lines indicate the parameter set that was found to be bistable after local stability analysis using eigenvalues of the Jacobian. D. Bifurcation diagram for the network in B showing the high and low stable states in solid lines and the saddle states in dashed lines. The shaded gray area is bistable. E. Deterministic simulation of the model with dashed gray lines indicating stable states. Stoichiometrically valid perturbations are provided every 0.2 s to the system and the system switches state at the third and the fourth perturbations. F. Stochastic simulation of the same model showing spontaneous transitions due to intrinsic thermal noise.

Each database entry is a model sampled from one of the reaction topologies, and includes an analysis of the three features described above. The entries are described using a unique hierarchical description ID which describes the model in this way: Class ID.Topology ID.Model ID. Thus the first model of the topology in **Fig 1B**, which has 3 reactants and 3 reactions is described as 3×3.113.1. The entry for each model is associated with the molecular concentration at steady-states, and the stability of each steady state (**Fig 1 B-E**). Additionally, we report final categories (singly stable, bistable, line, and other), the stable and saddle points, and the eigenvalues of the Jacobian matrix of the reaction rate system at each of these stable and saddle points. Each entry also includes a report on *stuck-states*. These are a special category of stable states relevant for stochastic chemistry (**Fig 1F**). The system can fall into such a state with a finite probability, but once in this state, it cannot transition out. For example, if A----> B, catalyzed by B, then there is a stuck state once B becomes zero.

This database serves a range of studies that examine stability in chemical systems and evolutionary analyses based on the relationships between reactions. For example (Siegal-Gaskins et al., 2009) have used a similar survey of possible model topologies to classify bistables in gene networks. We have recently used SWITCHES to study robustness in bistable systems (Moza and Bhalla, 2020). By providing searchable, crosslinked, and fully characterized models, the SWITCHES database allows for searches for features and relationships of such systems. Furthermore, by providing models in the standard SBML format, it enables access by over 100 software tools.

